# Explainable prediction of catalysing enzymes from reactions using multilayer perceptrons

**DOI:** 10.1101/2023.01.28.526009

**Authors:** Daniel Probst

## Abstract

Assigning or proposing a catalysing enzyme given a chemical or biochemical reaction is of great interest to life sciences and chemistry alike. The exploration and design of metabolic pathways or the challenge of finding more sustainable enzyme-catalysed alternatives to traditional organic reactions are just two examples of tasks that require an association between reaction and enzyme. However, given the lack of large and balanced annotated data sets of enzyme-catalysed reactions, assigning an enzyme to a reaction still relies on expert-curated rules and databases. Here, we present a data-driven explainable human-in-the-loop machine learning approach to support and ultimately automate the association of a catalysing enzyme with a given biochemical reaction. In addition, the proposed method is capable of predicting enzymes as candidate catalysts for arbitrary organic reactions. Finally, the introduced explainability and visualisation methods can easily be generalised to support other machine-learning approaches involving chemical and biochemical reactions.

---

The identification and classification of associations between enzymes and the reactions they catalyse is of great interest across fields. For biologists, this includes the mapping of an enzyme and its substrates into a metabolic network, which is an integral part of connecting experimental data with established domain knowledge, the creation of genome-scale metabolic models to enable the analysis of omics data, and the computational design of synthetic metabolic pathways (1–4). For medicinal chemists, the association of enzymes with their substrates is essential to the process of target-based drug discovery and to predict the metabolic fate of a drug candidate in an organism or specific organ (5, 6). Meanwhile, process chemistry and material science are interested in the discovery and engineering of enzymes that catalyse known or novel chemical reactions to produce new materials or drugs, increase the efficiency of synthetic routes, or replace existing synthetic routes with more environmentally friendly enzyme-catalysed alternatives (7–9). To support these efforts, a multitude of computational models have recently been proposed that enable the prediction of a required enzyme for a given reaction—generally relying on complex deep neural network architectures or expert-curated rules, both presupposing the existence of large data sets of biochemical reactions annotated with correct enzyme classifications (10–14). However, compared to other data in biology and chemistry, data sets containing annotated enzyme-catalysed reactions are exceedingly small and imbalanced in regard to enzyme-class distribution, complicating the application of data-hungry deep learning techniques, ultimately resulting in low predictive power—especially for underrepresented enzyme classes (12, 15–17).

The classification of enzymes and enzyme-catalysed reactions, and therefore their association, using EC numbers (see Methods for a detailed description of the EC number classification scheme) remains a manual task carried out by the Nomenclature Committee of the International Union of Biochemistry and Molecular Biology (IUBMB) (18, 19). With Rhea, the expert-curated knowledgebase of chemical and transport reactions of biological interest, an effort was started to annotate enzyme-catalysed reactions catalysed by enzymes found in UniProtKB with ChEBI identifiers and EC numbers (17). While this effort started to provide much-needed additional data, the requirement for extensive involvement of experts in the curation and classification of enzyme–reaction associations causes slow growth of annotated enzyme-catalysed reaction data sets. The resulting lack of data shows a need for automation in order to enable further progress in machine learning involving enzyme-catalysed reactions so as to reach the impact and utility of comparable approaches such as recent advancements in computer-assisted synthesis planning (CASP) that can draw from data sets containing millions of organic reactions (20, 21).

The available methods for the automated, computational classification of enzymes and their association with reactions can be divided into two approaches: Methods predicting an enzyme classification from the amino acid sequence or tertiary structure of an enzyme (22–25), and methods predicting an enzyme classification from the reaction catalysed by an enzyme, where the enzyme classification is the aforementioned EC number. The focus of this article is on the latter of the two approaches, assuming no knowledge of an enzyme’s sequence or structure. Many of the methods to computationally predict the class of an enzyme based on the catalysed reaction rely on the explicit mapping or typing of atoms, bonds, or functional groups; a predefined set of physicochemical and topological descriptors; or the balancing of reaction equations, which requires significant preprocessing and manual curation (26–31). Other approaches include similarity searches based on molecular fingerprints and substructure matching algorithms (32, 33). While the average accuracy of these data-driven methods has been high, they remain error-prone in edge cases and in predicting the many enzyme subclasses and sub-subclasses where little training data is available. A notable recent approach, setting the state-of-the-art, is the rule-based approach BridgIT by Hadadi et al. (34). However, while this method has excellent accuracy and allows for explainability due to its rule-based rather than data-driven nature, the rules have to be created and continuously updated by experts with a deep knowledge of enzyme-catalysed reactions. This superior performance of rule-based compared to data-driven methods, the continuing need for expert curation of databases, and the existence of commission oversight show a need that goes beyond current approaches (17). While the superior performance of rule-based methods can be attributed to a lack of sufficiently large and balanced data sets, the lack of adaption of an automated annotation process can have multiple possible reasons, including the absence of model explainability or trust, a lack of utility, or a failure to identify the limits of applicability (35, 36). A potent solution to these is explainable machine learning, which can increase acceptance and usability as well as identify the limits and edge cases of a model, and has been widely used in genetics, healthcare, or education (37–39). However, recent approaches in explainable machine learning in chemistry remain limited to models trained on single molecular entities rather than reactions (40–42).

Here, we introduce explainability to a multilayer perceptron that predicts the EC number of an enzyme given a reaction without the need for balancing the reaction or any other form of reaction curation. We provide a tool that can support and eventually fully automate enzyme–reaction association and classification by introducing three main advancements. (i) We report multiple models capable of predicting the classes, subclasses, and sub-subclasses of enzymes that catalyse a given reaction with overall accuracies of 98, 97, and 95 per cent, respectively, while requiring minimal training resources, enabling continuous retraining. (ii) By mapping the molecular fragments occurring in the reactions to the vector entries that act as input for a neural network classifier, we enable the use of a modified version of the DeepLIFT algorithm (DeepSHAP) to annotate fragments and atoms with their respective classification contributions, providing chemical explainability for all described models. (iii) We develop and implement a generalised approach for the visualisation of numerical annotations for molecules and reactions, which we use to visualise the classification contributions that explain a model’s perception of an input reaction. Based on these advancements, we introduce an approach that can be used as a human-in-the-loop machine-learning solution for the transition to the fully automated annotation of enzyme-catalysed reactions. Furthermore, our system allows for the prediction of catalysing enzyme candidates for arbitrary reactions, making it a utility for the exploration of the enzymatic reaction space by chemists and biologists alike. The resulting models, data, and libraries are made accessible as a hosted web application and a locally installable Python package that includes a graphical and command-line interface in addition to a Python API. The modular architecture of the system allows for easy extension or the reuse of specific components in virtually all machine-learning tasks involving chemical or biochemical reactions.

## Results

### Enzyme classification using differential reaction fingerprints and a simple multilayer perceptron

Enzyme-catalysed reactions are stored as reaction SMILES, a string representation of a chemical reaction based on the molecular graphs of the participant substances (43). As a first step, we encode these string representations into a binary vector using the differential reaction fingerprint (DRFP), which we recently showed to provide state-of-the-art reaction representations by example of reaction yield predictions, performing at least as well as DFT-derived descriptors or transformer-based methods on a yield prediction task for organic reactions (44). A TMAP visualisation shown in 2 exemplifies the ability of DRFP-encoded enzyme-catalysed reactions, extracted from Rhea (*n* = 7, 010), to be classified using the EC numbering scheme. Figure 2a is a visualisation of the entirety of the Rhea database coloured by the enzyme classes oxidoreductases (EC 1), transferases (EC 2), hydrolases (EC 3), lyases (EC 4), isomerases (EC 5), and ligases (EC 6). Figure 2b is a detailed view, displaying only transferases, coloured by the nine transferase subclasses found in Rhea: EC 2.1 (transferring one-carbon groups), EC 2.2 (transferring aldehyde or ketonic groups), EC 2.3 (acyltransferases), EC 2.4 (glycosyltransferases), EC 2.5 (transferring alkyl or aryl groups, with the exception of methyl groups), EC 2.6 (transferring nitrogenous groups), EC (transperring phosphorus-containing groups), EC 2.8 (transferring sulfur-containing groups), and EC 2.9 (transferring selenium-containing groups). Finally, Figure 2c shows the three sub-subclasses of glycosyltransferases, namely, hexosyltransferases (EC 2.4.1), pentosyltransferases (EC 2.4.2), and those transferring other glycosyl groups (EC 2.4.99). As these plots illustrate, DRFP is able to separate reactions according to all three levels of the EC classification. Furthermore, the plot shows that the fingerprint is capable of separating oxidoreductases (EC 1) exceedingly well, while there is a relative lack of distinct clustering for isomerases (EC 5). These findings reflect previous observations and are primarily caused by the diversity of reactions within a class (12).

**Fig. 1.**
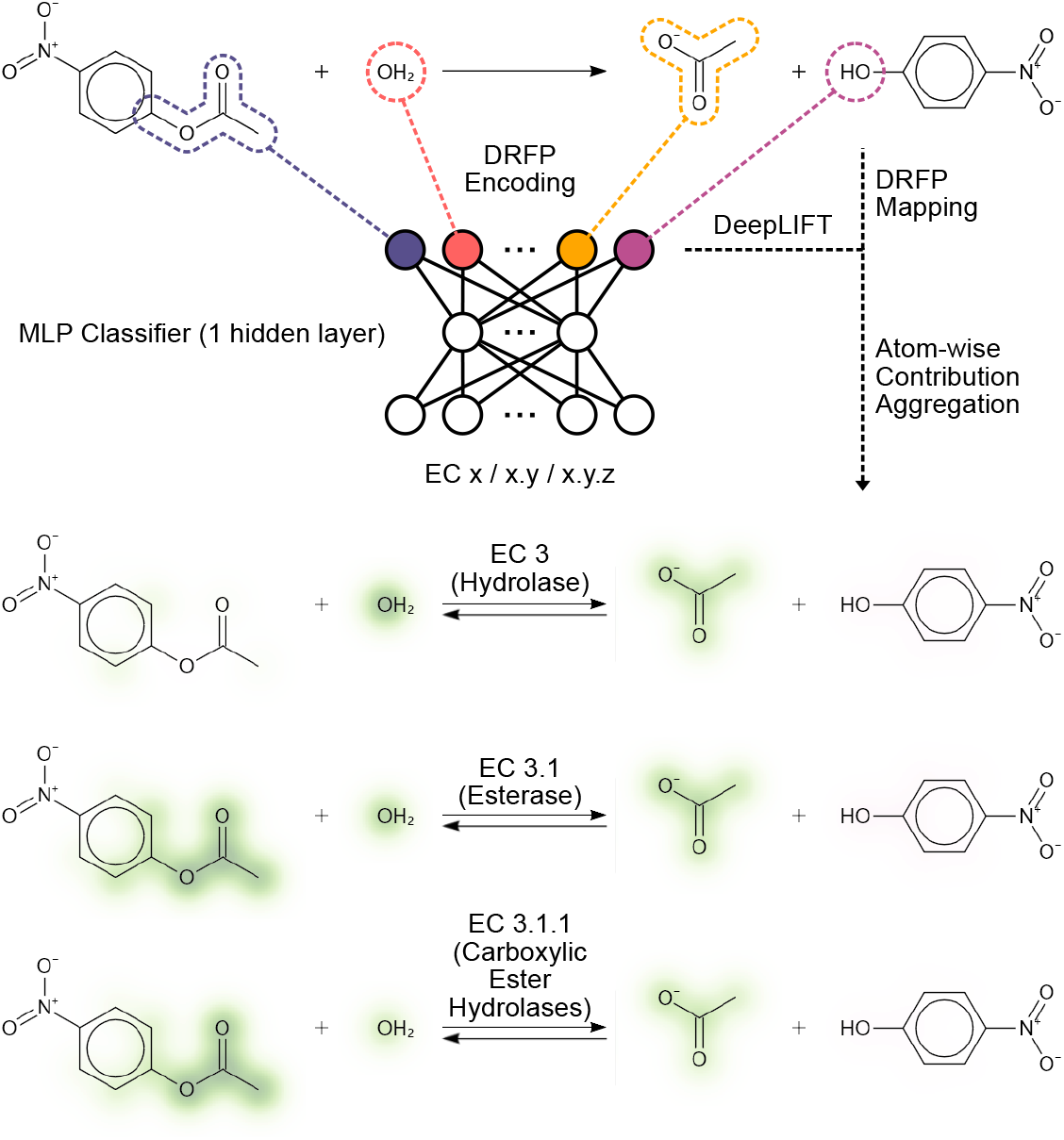
Explaining enzyme-catalysed reaction classifications. Enzyme-catalysed reactions are classified using three different models. Model ECX predicts the class, model ECXY predicts the class and subclass, and model ECXYZ predicts the class, the subclass, and the sub-subclass. The input to all models is a binary DRFP encoded reaction SMILES that allows a mapping between input and molecular fragment. Using Shapley additive explanations (SHAP), the inputs’ influence on the classification can be traced back to a molecular fragment.

**Fig. 2.**
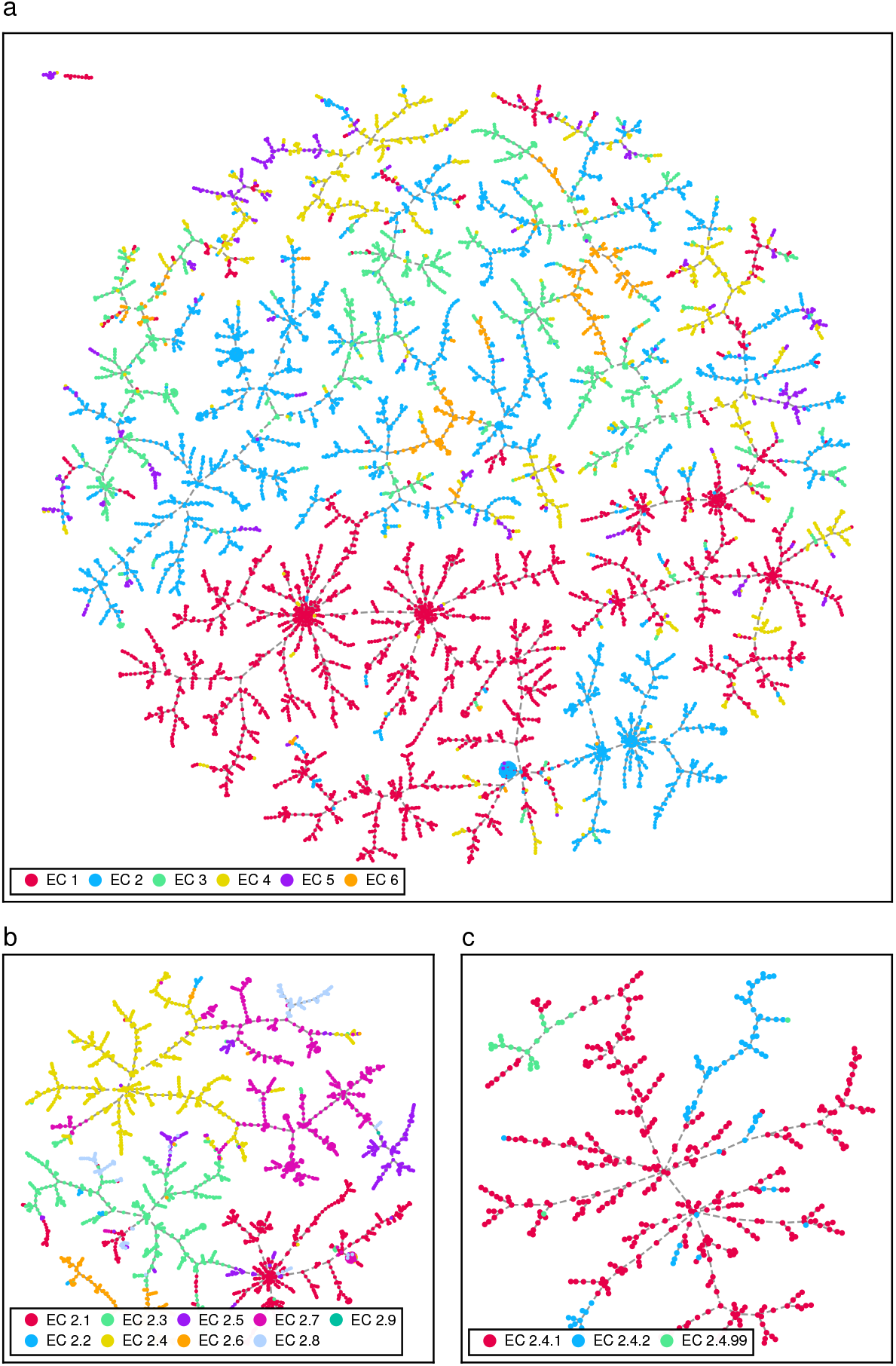
TMAPs of DRFP-encoded biochemical reactions extracted from the Rhea database coloured by the associated EC number. (a) All reactions coloured by enzyme class. (b) Transferase-catalysed reactions coloured by subclass. (c) Glycosyltransferase-catalysed reactions coloured by sub-subclass.

Following the encoding of the enzyme-catalysed reactions extracted from Rhea as DRFP fingerprints, three distinct models were trained on the data: ECX, ECXY, and ECXYZ. While all models share a simple multilayer perceptron (MLP) architecture with a single hidden layer, ECX, ECXY, and ECXYZ were trained with labels representing x.-.- (classes), x.y.- (classes and subclasses), and x.y.z (classes, subclasses, and sub-subclasses), respectively. The specific architecture of the MLPs is described in Methods. In addition to the Rhea-extracted data, the procedure was repeated for our recently released ECREACT data set (*n* = 81, 205), which extends the reactions from Rhea with reactions extracted from BRENDA, PathBank, and MetaNetX (12, 45–47). The accuracies and f-scores of the models are shown in Table 1 together with the training times as well as the training and experimentation energy use. Figure 3 shows the confusion matrices for the tests of ECX_Rhea_, ECXY_Rhea_, and ECXYZ_Rhea_ in the first row (a-c) and the confusion matrices for ECX_ECREACT_, ECXY_ECREACT_, and ECXYZ_ECREACT_ in the second row (d-f). Given the different sizes of the classes, subclasses, and sub-subclasses, the boxplots in Supplementary Figure S2 yield further insights into the existence of challenging, low-accuracy cases of subclasses and sub-subclasses that have little effect on the overall accuracy when being evaluated together with the larger, better-trained classes but represent important edge-cases. For both Rhea and ECREACT, these challenging cases resulting in low prediction accuracies are generally caused by (sub-)subclasses with a small number of samples, rather than larger (sub-)subclasses with diverse samples (Supplementary Figures S3 and S4). This behaviour follows the examples established in our previous work on biocatalysed synthesis planning (12). The comparatively poor overall accuracy of isomerases (EC 5) can be explained by their function, which is to carry out modifications within a molecule, that would be assigned to other enzyme classes if they took place between two different molecules. This is shown in Figure 3a,b where potential intramolecular transferases, which are classified as isomerases (EC 5), have been classified as intermolecular transferases (EC 2). Overall, the observations show that a lack of training samples in under-represented classes, subclasses, and sub-subclasses leads to low accuracies in data-driven machine-learning approaches. The following sections will introduce the methods and tools necessary to facilitate the bridging of expert curation and data-driven learning of enzyme-catalysed reactions to speed up data curation and explain a model’s behaviour to scientists.

**Table 1.** Accuracies and F-scores of the different models trained on Rhea and ECREACT. The energy use is calculated based on the energy use of the device (Dell XPS 15, i7-12700H CPU, NVIDIA GeForce RTX 3050 Ti Laptop GPU) and includes the power usage of models trained for 4x cross-validation and four experiments with fingerprint variations. The hyperparameter values were taken from the previous work on organic reactions (44). The total resulting energy consumption for model experimentation, training, and validation for this project was 3.25 kWh. Energy mix (2022): 65% hydro, 23% solar, and 12% other renewables.

| Model | Accuracy | F-Score | Training Time | Energy Use |
| --- | --- | --- | --- | --- |
| ECX<sub>Rhea</sub> | 0.98 ± 0.00 | 0.97 ± 0.01 | 100 s | 43 Wh |
| ECXY<sub>Rhea</sub> | 0.96 ± 0.01 | 0.88 ± 0.02 | 160 s | 70 Wh |
| ECXYZ<sub>Rhea</sub> | 0.95 ± 0.00 | 0.87 ± 0.02 | 190 s | 82 Wh |
| ECX<sub>ECREACT</sub> | 0.98 ± 0.00 | 0.96 ± 0.00 | 2,090 s | 904 Wh |
| ECXY<sub>ECREACT</sub> | 0.95 ± 0.00 | 0.82 ± 0.01 | 2,170 s | 936 Wh |
| ECXYZ<sub>ECREACT</sub> | 0.93 ± 0.00 | 0.77 ± 0.01 | 2,810 s | 1,216 Wh |

**Fig. 3.**
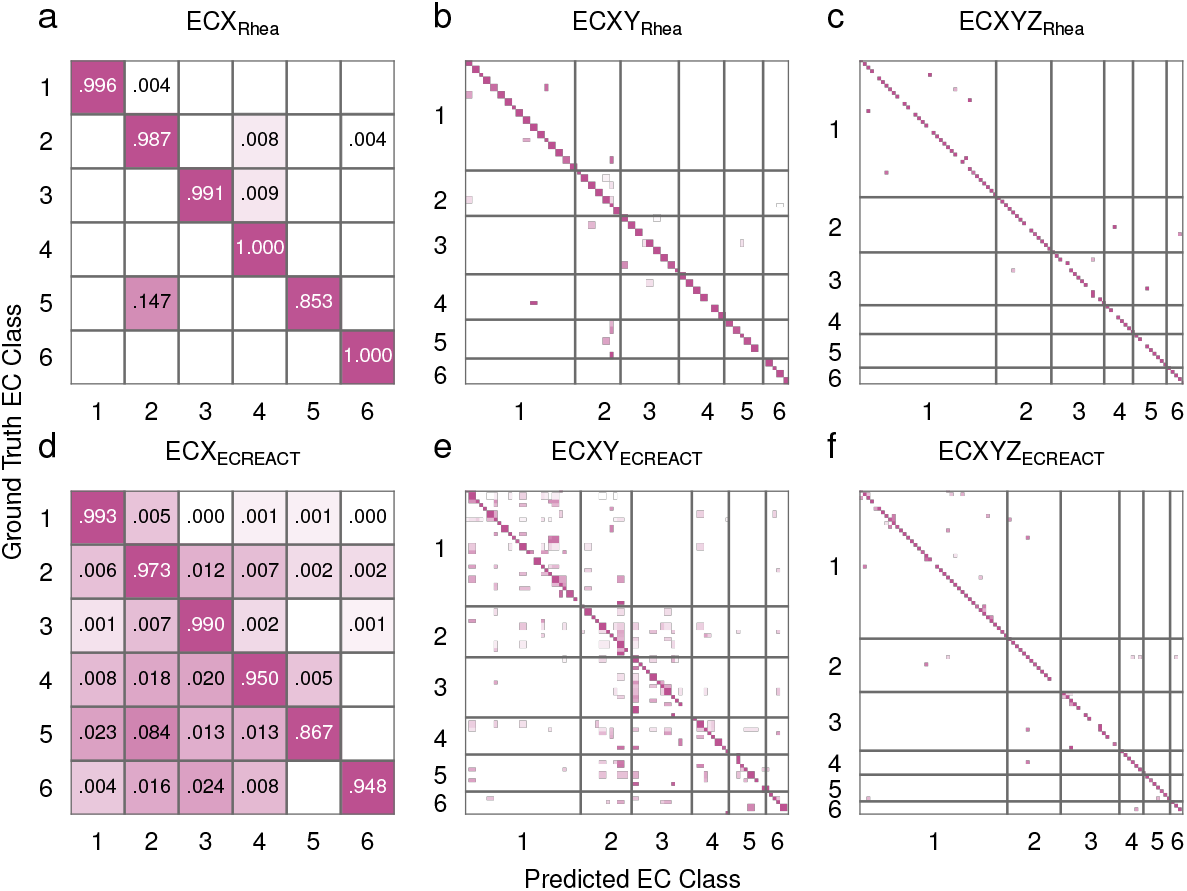
Confusion matrices for reaction-based enzyme classification. (a,d) Enzyme class-level confusion matrices for models trained on the Rhea and ECREACT data sets, respectively. (b,e) Subclass-level confusion matrices for models trained on the Rhea and ECREACT data sets, respectively. (c,f) Sub-subclass-level confusion matrices for models trained on the Rhea and ECREACT data sets, respectively.

### Explaining classification using DeepLIFT

Using the differential reaction fingerprint to embed the reaction as a binary vector for input into the MLP allows for a mapping between binary input feature *v*_*i*_ and molecular fragment *f*_*i*_. This enables the use of an arbitrary approach for explainable machine learning capable of determining or estimating the contribution of an input feature *v*_*i*_ to a resulting classification, to also quantify the influence of each molecular fragment *f*_*i*_. Based on its ease of use and remarkable performance, we selected the DeepSHAP implementation of DeepLift to estimate the contributions of input features to the classification (48, 49). Given the fragment contributions 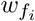, the atom contributions 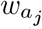 are calculated by summing 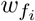 of a given reaction (*v*_*i*_ = 1) that include atom *a*_*j*_.

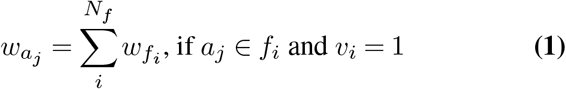

This atom-wise weighing enables later visualisation of overlapping or contained fragments. The result of this operation can be seen in Figure 4, where fragments with positive weights that contribute towards a certain classification are coloured green, while fragments with negative weights that contributed against a certain classification are shown in magenta. However, not only fragments present in the reaction can have an effect on the classification. The absence of a certain fragment can influence the classification as much as the presence of another. Therefore, the information on the contributions of absent fragments is retained to provide a more complete picture of the model’s decision at a later point (Figure 5).

**Fig. 4.**
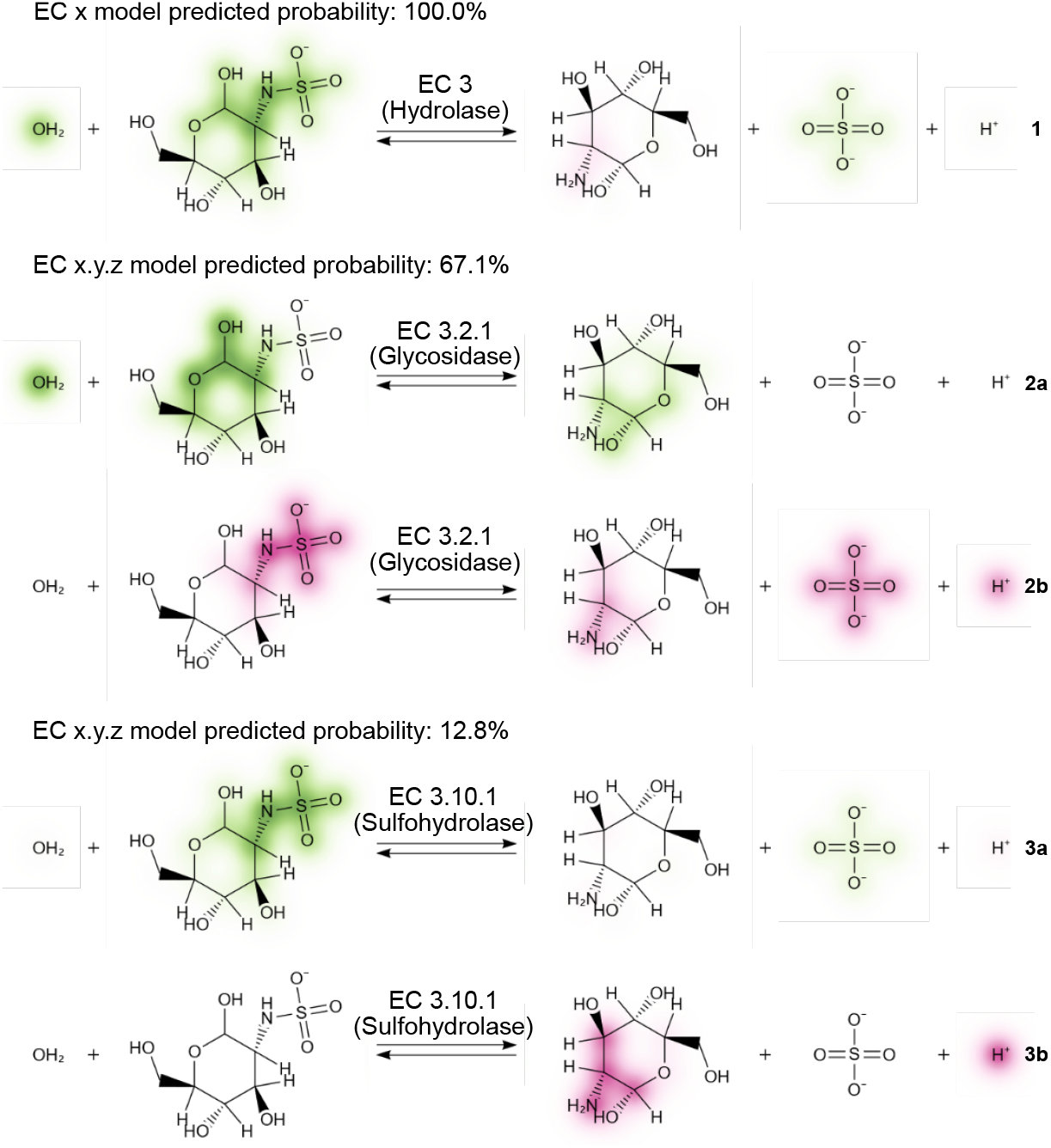
Explaining a misclassification of a N-sulfoglucosamine sulfohydrolase (EC 3.10.1) as a glycosidase (EC 3.2.1). The model ECXRhea correctly predicts the reaction (**1**) to be catalysed by a hydrolase (EC 3), primarily focusing on the water (OH2) and the hydrolysed bond, both with a positive contribution towards EC 3. In addition, there is a small negative contribution against EC3 shown on the amine group. Unlike **1**, where positive and negative contributions are shown in one reaction drawing, positive and negative contributions are split into separate depictions for **2** and **3** for visualization purposes. For the top prediction (67.1%) of model ECXYZRhea the focus of the model shifts to a non-reactive site including a hydroxy group in the N-sulfo-D-glucosamine as a major positive contribution (**2a**), while the sulfur-nitrogen bond is the major negative contribution (**2b**). For the correct prediction (top-2, 12.8%), the model remains focused on the hydrolised sulfur-nitrogen bond with a positive contribution (**3a**) as the negative contributions (**3b**) can be found on the D-glucosamine and the proton.

**Fig. 5.**
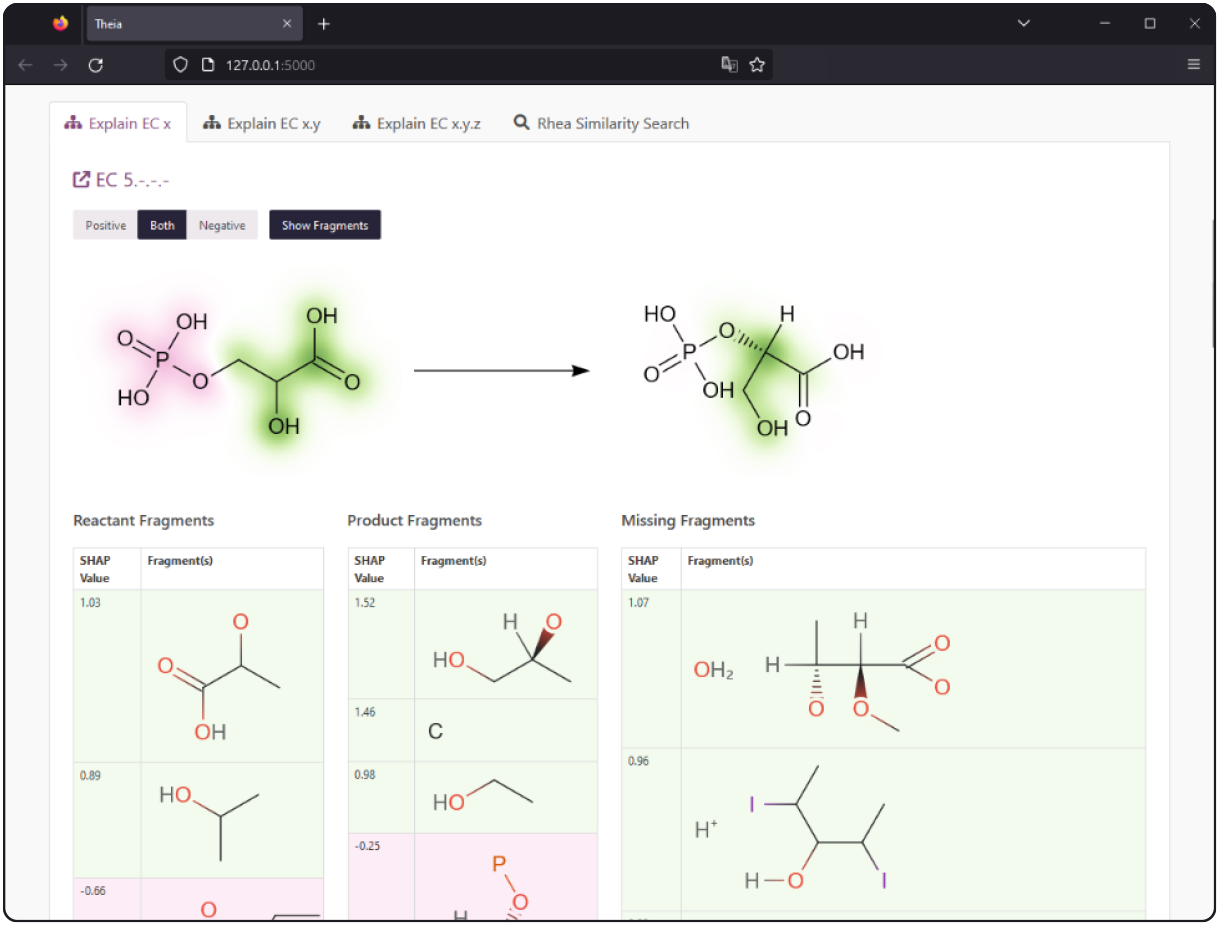
Visualising the contributions to the classification of a reaction to being catalysed by an isomerase (phosphoglycerate mutase, EC 5.4.2) using a web interface. Fragments that are not present in a participating molecule yet still contribute heavily towards the predicted class are displayed with their respective contribution. In addition, for entries in the binary input vector that represent multiple fragments, all the associated structures are represented.

A potential caveat in mapping the contribution values to a fragment is collisions. Collisions can happen at two stages of the method: (i) When hashing the SMILES representation of the sub-structures from a reaction to a set of 32-bit integers and (ii) when folding the 32-bit integers into a fixed-size binary vector using a modulo operation. The number of collisions of a 32-bit hashing function can be estimated based on the maximum hash value and the number of unique fragments using a generalisation of the birthday problem (50). For a maximum hash value of 2^32^− 1 and 9,509 and 16,983 unique fragments extracted from Rhea and ECREACT,respectively, this results in 0.01 expected hash collisions for Rhea and 0.03 for ECREACT. However, when folding the sets of 32-bit integers into 10,240-dimensional binary vectors using a modulo operation, 2,439 and 5,060 of the entries represent more than one fragment for Rhea and ECREACT, respectively. While the models still perform well given these collisions, as shown by our previous work (44, 50), this could potentially negatively impact the interpretability of the model using DeepLIFT values, especially if multiple fragments of a given reaction are represented by the same entry in the binary vector. For the visualisation of fragments that are part of a given reaction, this is a minor concern, as two fragments occupying the same entry only occur 147 times (2.1 %) in Rhea (*n* = 7, 010) and 2,025 times (2.5 %) in ECREACT (*n* = 81, 205). For fragments that are not part of a given reaction yet still contribute to the decision of the model, however, this is not the case. We solve this problem and introduce a generalisable approach to the visualisation of explainable machine learning for reactions in the following section.

### Visualising explanations using reaction depictions

The visualisation of the classification contributions calculated using the DeepSHAP implementation of DeepLIFT as shown in Figure 4 is enabled by implementing an upgrade to the previously released SmilesDrawer JavaScript library (51). The script takes the reaction SMILES and the atom contributions as computed in Equation 1 as an input. As a first step, the atom contribution values 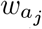 are normalised across all molecules in the reaction to show the relative contributions. Next, the contributions 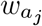 are assigned the coordinates of the respective atoms *v*_*j*_ = 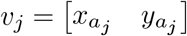 1 in each molecules 2D drawing space. Finally, the colour values for the pixel grid are assigned by the summatory function *G* of all bivariate gaussian distributions centred at *v*_*j*_ with an arbitrary *σ*.

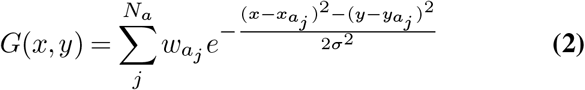

Choosing a diverging colour scale to represent the contribution values at each coordinate in the pixel grid produces an intuitive representation of the contribution values. The resulting visualisation can be seen in Figure 4. However, this only enables the visualisation of the contributions of fragments that are present in a given reaction. In order to also visualise the contribution of fragments that are not present in a reaction, these fragments are listed with their respective value as part of a web service or a locally run program. Figure 5 shows an example of a phosphoglycerate mutase (EC 5) catalysed reaction as explained in the web application on which the introduced approach is being made available. The contributions towards being classified as an isomerase come as much from the absent fragments, a proton (H^+^) and water (OH_2_), as from the fragments found in the participating molecules. The example of the interface shown in Figure 5 also introduces how the occurrence of multiple fragments assigned to the same entry of the binary input vector can be handled by displaying the colliding substructures. In this case, it is trivial to determine the influential missing fragments, water and proton, from the context of the given reaction and the predicted class (EC 5).

To facilitate access to the presented visualisations, the presented models are deployed as a web application and a Jupyter notebook, as well as graphical and command line interfaces installable via Python’s pip package manager.

## Discussion

The approach presented in this work introduces a way forward in enzyme–reaction classification beyond expert curation. The introduced models and software will initially support the growth and balancing of databases containing annotated enzyme-catalysed reactions such as Rhea through human-in-the-loop machine learning. The utility of this approach is illustrated in Figure 4, where the explanation of a misclassification of the model lends insight into the underlying causes of the inaccuracy such as a lack of training data of certain classes, in this case, Sulfohydrolase-catalysed reactions. Based on such information, an expert curator can modify either the architecture of the model or the composition of the training data set. The fast and efficient training of the classifiers (below 10 minutes using approximately 25 Wh on a consumer laptop) then allows for continuous retraining with an adjusted architecture or on newly annotated or balanced data. Eventually, as the classes with sufficient examples show, the presented solution will be able to take over reaction-based enzyme and enzyme function classification from humans. In their current iteration, the trained models can already be used to predict a candidate catalysing enzyme for an arbitrary chemical reaction, while the software allows for easy human evaluation of the predictions. In addition, the generalisable method to visualise explainable machine learning for chemical and biochemical reactions can be adapted by current and future machine learning tools involving arbitrary explainability techniques and machine learning architectures. Documentation, code, and notebooks showcasing the described functionality as well as instruction to install the tools locally from PyPI can be found at https://github.com/daenuprobst/theia.

## Methods

### EC Classification

Once identified and associated with a biochemical reaction through experimentation, enzymes are classified according to their function and the reactions they catalyse using the hierarchical EC Number (Enzyme Commission Number) scheme based on the reaction they catalyse. This hierarchical classifier, in the form x.y.z.sn, where x is the class, y is the subclass, z is the sub-subclass, and sn is an incremental serial number assigned to an enzyme. The class (x) encompasses seven categories: (1) Oxidoreductases, (2) Transferases, (3) Hydrolases, (4) Lyases, (5) Isomerases, (6) Ligases, and (7) Translocases. While classes 1 through 6 catalyse a chemical modification of the substrate, translocases are limited to catalyse the movement of molecules or ions across membranes. Translocases are, therefore, not within the scope of this study. The subclass (y) of an enzyme specifies the group or bond on which the enzyme acts. For example, hydrolases of subclasses 3.4 and 3.7 act on peptide and carbon-carbon bonds, respectively. The sub-subclass (z) of an enzyme further specifies the reaction. Peptidases (3.4) with the sub-subclass 3.4.13 are dipeptidases with dipeptides as a substrate, while peptidases with the sub-subclass 3.4.22 are cysteine endopeptidases that hydrolyse peptide bonds after non-terminal cysteines. Finally, the serial number (sn) does not convey information on the reaction but distinguishes different enzymes that catalyse the same types of reactions.

### Data processing

The ECREACT data set does not require any preprocessing as it is available as a `.csv` file containing reaction SMILES and the associated EC number. For Rhea, the file containing the reactions annotated with ChEBI identifiers is downloaded and then processed with a ChEBI export to match the molecular identifiers with the respective SMILES to generate reaction SMILES. The processed data, as well as a shell script to download the required data and a Python script to process the raw Rhea data are included in the GitHub repository.

### Differential reaction fingerprint (DRFP)

For encoding the reaction SMILES as differential reaction fingerprints using the drfp PyPI package (version 0.3.6), the parameters were chosen to maximise accuracy while minimising the probability of collisions, in order to enhance explainability. The folded length (dimensionality) of the DRFP fingerprint was chosen as 10,240 (default 2,048) and the radius as 2 (default 3). The DRFP encoding function was adapted to produce non-centred canonicalised SMILES, further reducing the number of potential collisions. Whereas the original function would produce multiple SMILES rooted at each atom for each fragment (e.g. `COC, OCC`, and `CCO` for dimethyl ether), the adapted version produces only a single SMILES per fragment (e.g. COC for dimethyl ether). This change can be toggled in the updated DRFP package using the argument `root_central_atom` in the static function encode. Furthermore, the function was adapted to explicitly include hydrogens in the SMILES encoding of the fragments (e.g. `[H]C([H])([H])OC([H])([H])[H]` instead of `COC`). This change can also be toggled in the updated DRFP package using the argument `include_hydrogens` in the static function encode.

### Multilayer perceptrons

The multilayer perceptrons used for all models in this work were implemented using PyTorch (version 1.13.0). The initial hyperparameters were taken from our previous publication on DRFP (44). The MLP consists of a linear input layer with 10,240 nodes, a hidden layer with 1,664 nodes, and a linear output layer with a number of nodes that is equal to the number of classes (unique EC numbers). Cross entropy with default parameters is chosen as the loss function (criterion), and Adam with a learning rate of 0.001 as optimiser. PyTorch’s exponential learning rate scheduler with a gamma of 0.9 is set as the scheduler. Finally, early stopping is implemented by monitoring the mean validation loss of the 5 most recent epochs. The training is stopped if the improvement of the current loss drops below 1.1. Training and validation losses for all models are shown in Supplementary Figure S1.

### DeepSHAP explanations

DeepSHAP is an extension of DeepLIFT, an additive feature attribution method, based on the assumption that DeepLIFT approximates SHAP values. A detailed description of the method can be found in the section *Deep SHAP (DeepLIFT + Shapley values)* of Lundberg and Lee (49). Using the described method, DeepSHAP assigns each feature (molecular fragment) an importance value for a given prediction based on a baseline value. The baseline value is calculated from a set of samples—100 reaction SMILES in the presented implementation—and represents an approximation of the average of all predictions. The SHAP (SHapley Additive exPlanation) values that measure the contributions of a feature based on the baseline are then the summed Shapley values of a conditional expectation function of the original model Lundberg and Lee (49).

### TMAP visualisation

The TMAP visualisations shown in Figure 2 were generated using the PyPI package tmap-viz. The parameters `sl_repeats=2, mmm_repeats=2, and n_trees=50` remained constant for all three subfigures, while while `node_size` was set to 3, 2, and 10 for subfigure a, b, and c, respectively. The script to generate the TMAPs is available in the project’s GitHub repository.

### Reaction visualisation

The reaction visualisations are based on the SmilesDrawer JavaScript library that, compared to other available libraries, allows the depictions of molecules and reactions in web applications without the need for server-side image rendering (51). To enable the visualisation of numeric attributes on a per-atom level, the library was extended with the ability to draw arbitrary pixel values on a background layer. While the reactions can be rendered as rasterised images (HTML canvas, or image elements) or vector images (HTML SVG elements), the background layer is always rendered as a rasterised image and scaled without interpolation for performance reasons. Finally, a wrapper for the display of explainable reactions with the SmilesDrawer JavaScript library in Jupyter notebooks is available on PyPI in the package `faerun-notebook`.

### Web application

In order to make the models and visualisations easily accessible, the presented approach is deployed as a Flask-based web application. In addition to the EC number predictions and the visualisation of the molecular contributions, the application also performs a nearest neighbour search of the DRFP-encoded reaction on the Rhea data using an Annoy index (52). In addition to the hosted version, the application is available as Docker and PyPI packages for on-premise deployment or local use.

## ACKNOWLEDGEMENTS

I would like to thank Prof. Pierre Vandergheynst for hosting me in his research group as well as the Swiss-Prot group at the Swiss Institute of Bioinformatics for their fruitful feedback and discussions.

## Data Availability

All used data sets are available as `.csv` files on GitHub. The training, validation, and test sets can be recreated using Python scripts available in the same GitHub repository https://github.com/daenuprobst/theia.

## Code Availability

All implementations and used software are available under open-source licences. The code for the web application as well as the scripts necessary to reproduce this work can be found at https://github.com/daenuprobst/theia. The repositories of the adapted DRFP and SmilesDrawer code are https://github.com/reymond-group/drfp and https://github.com/reymond-group/smilesDrawer, respectively.

## Supplementary Figures

**Fig. S1.**
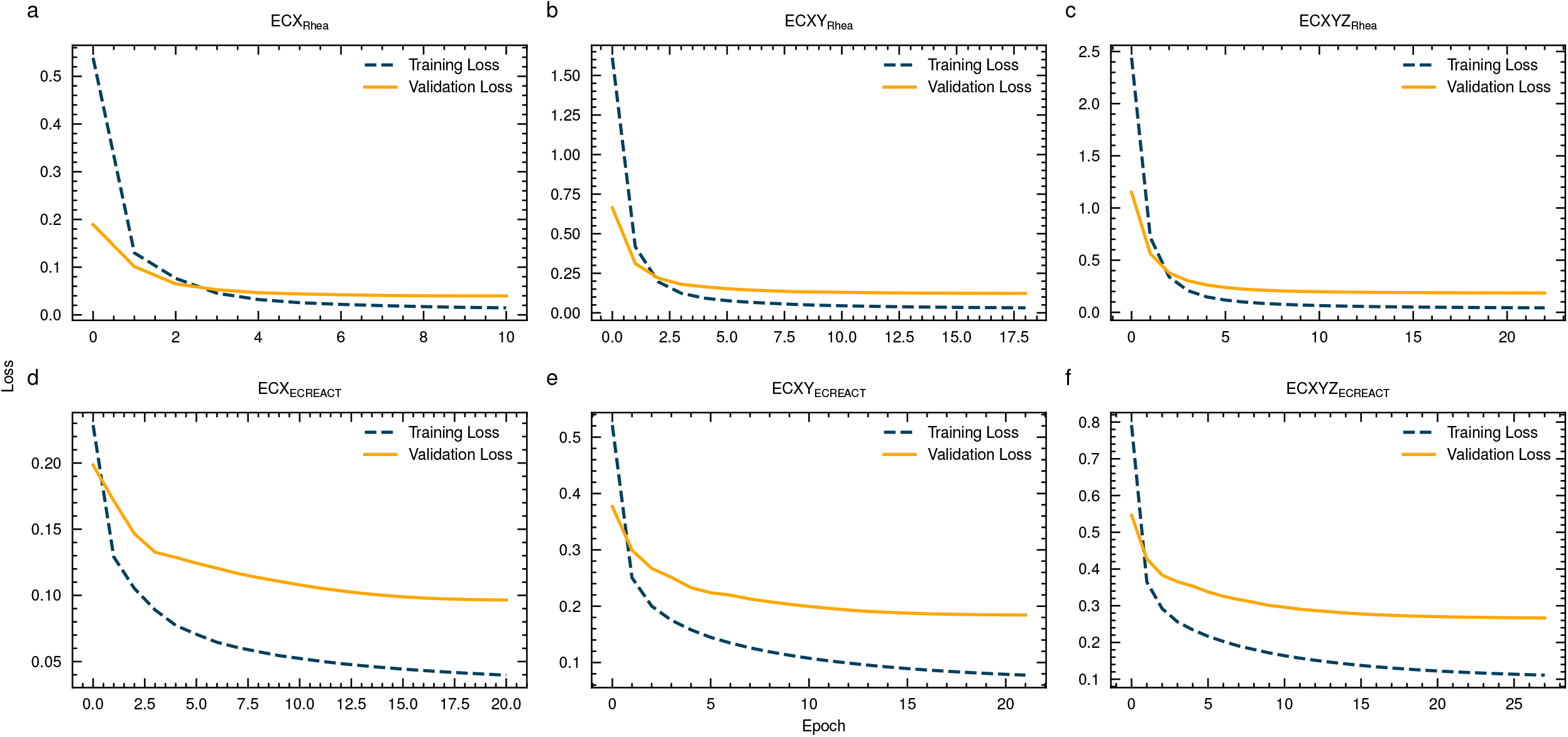
Training and validation losses for the models presented in this work. Early stopping is implemented by monitoring the mean validation loss of the 5 most recent epochs. The training is stopped if the improvement of the current loss drops below 0.001.

**Fig. S2.**
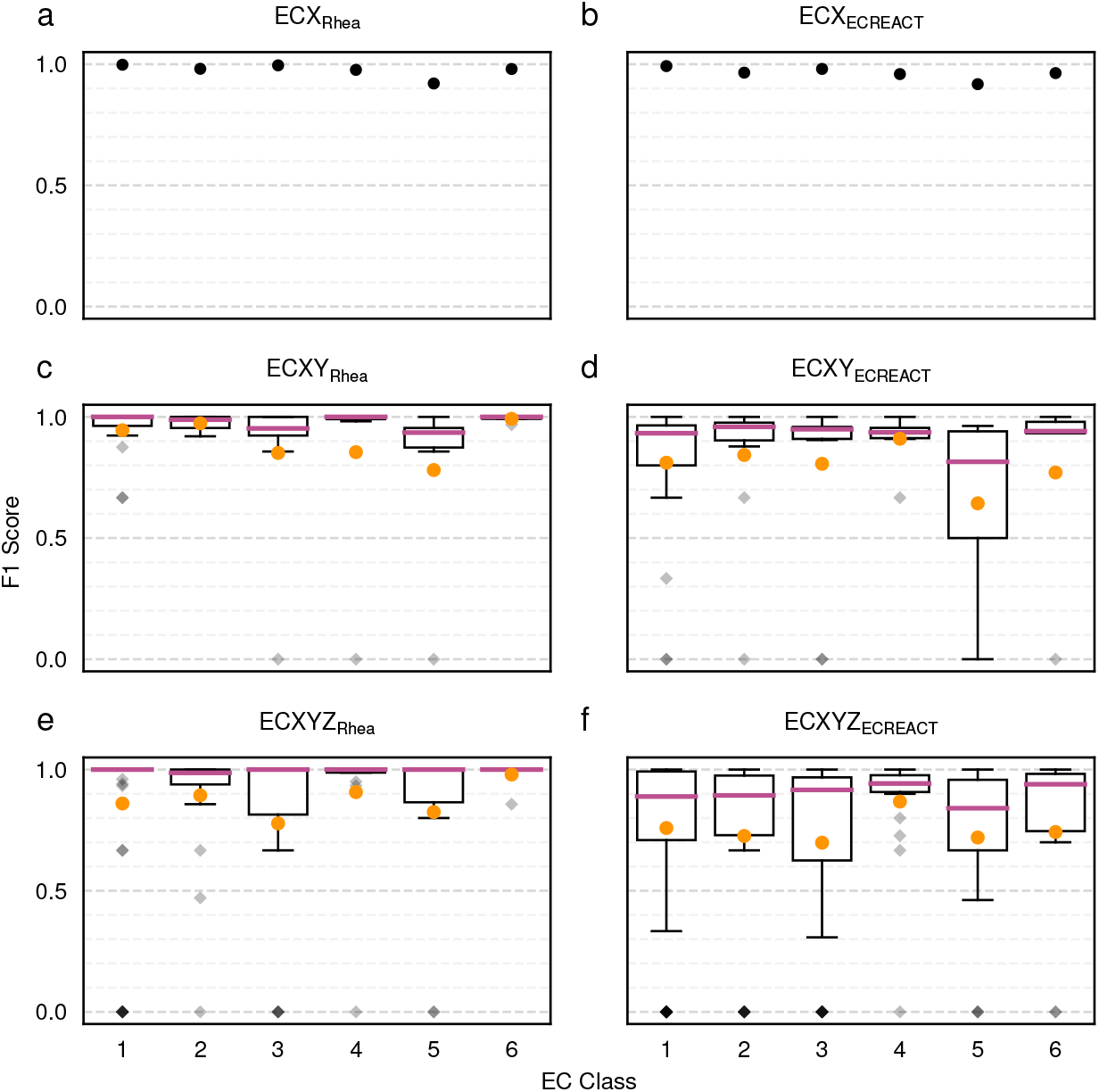
Boxplots showing the distribution of accuracies among classes, subclasses, and sub-subclasses. The filled orange circle represents the mean.

**Fig. S3.**
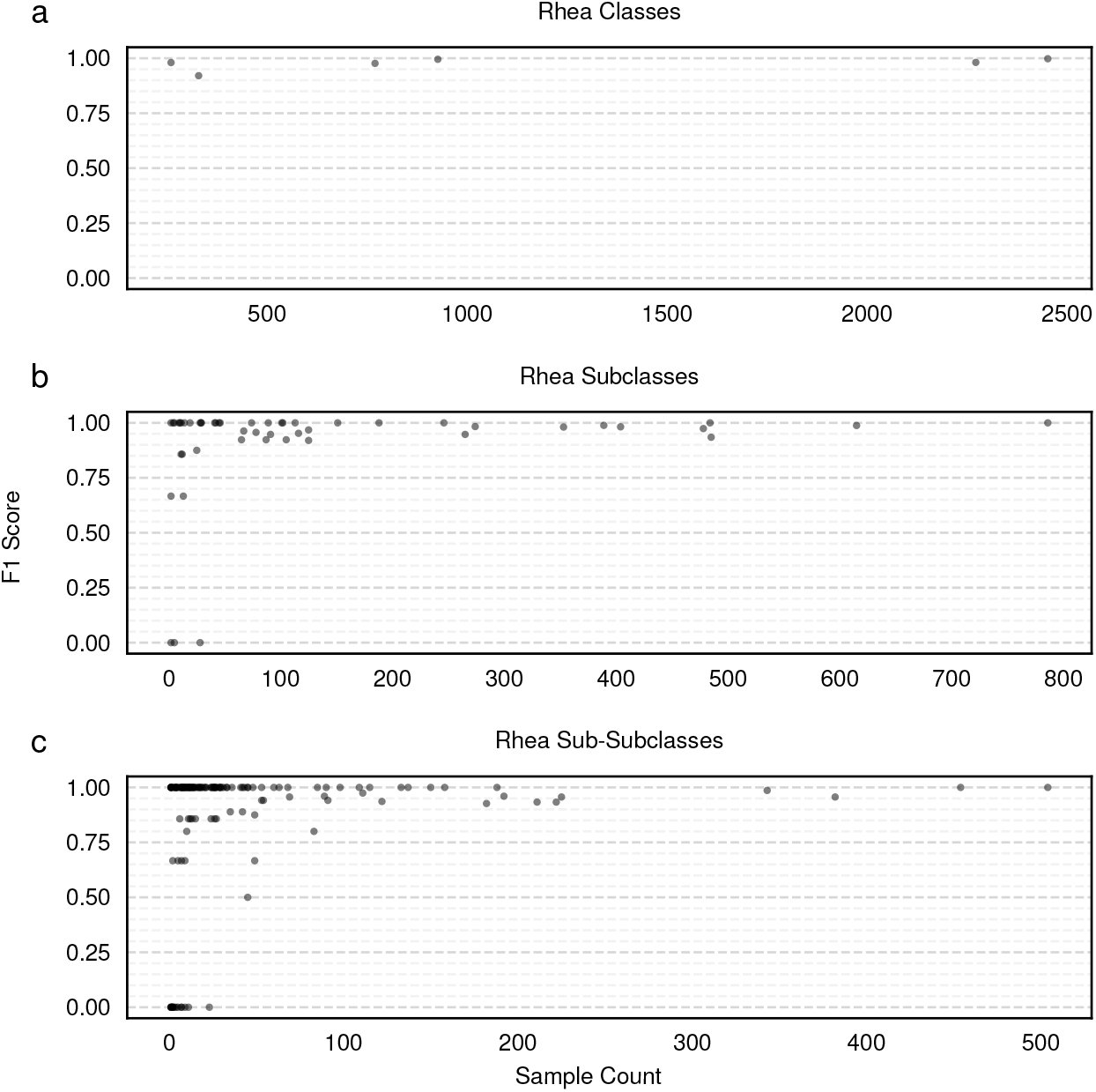
Scatter plots showing the dependence of accuracy on sample size for models trained on Rhea data.

**Fig. S4.**
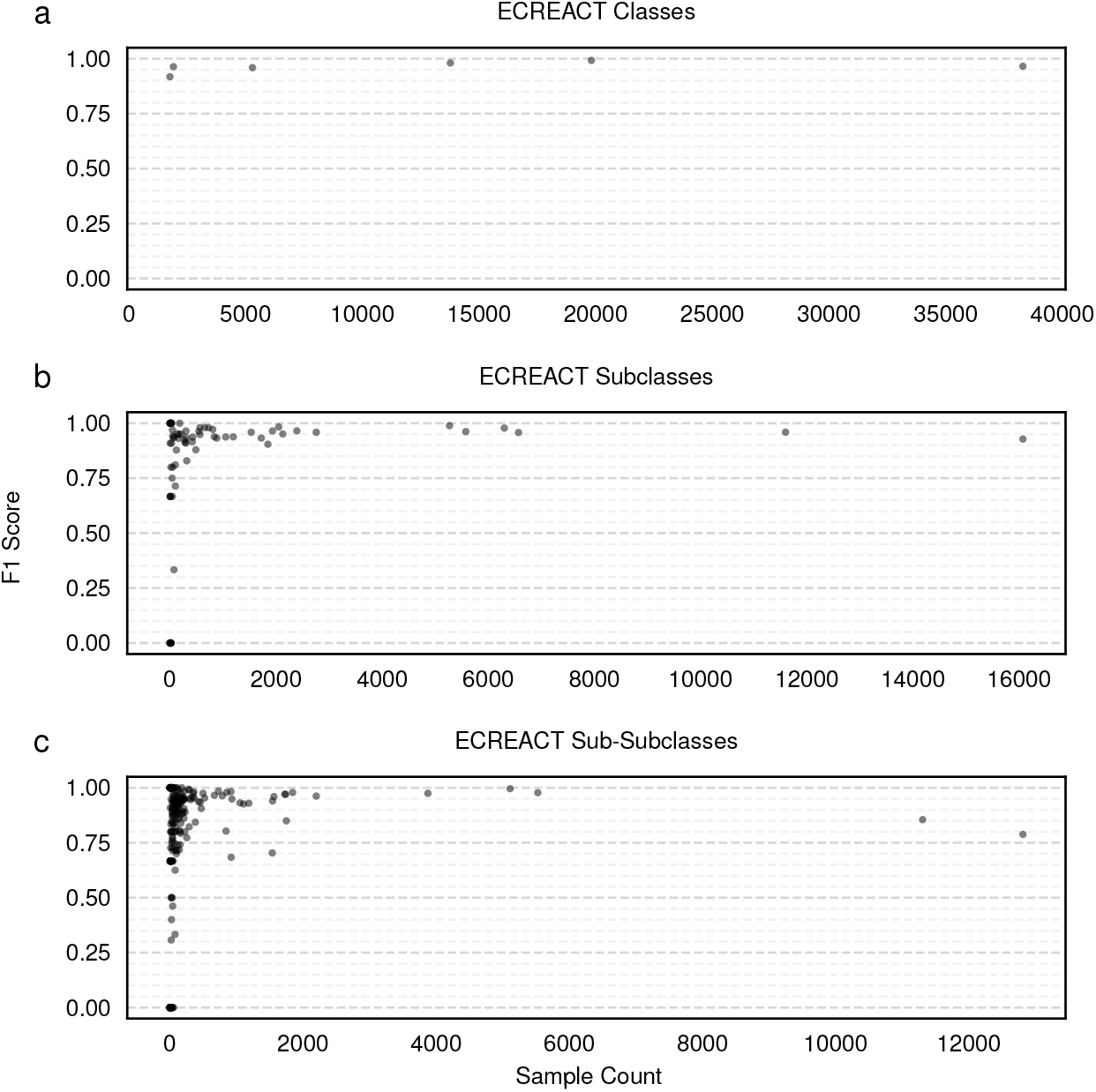
Scatter plots showing the dependence of accuracy on sample size for models trained on ECREACT data.

## References

1. Minoru Kanehisa, Miho Furumichi, Mao Tanabe, Yoko Sato, and Kanae Morishima. KEGG: new perspectives on genomes, pathways, diseases and drugs. Nucleic Acids Research, 45(D1):D353–D361, 2017. ISSN 0305-1048. doi: 10.1093/nar/gkw1092.

2. D.-S. Lee, J. Park, K. A. Kay, N. A. Christakis, Z. N. Oltvai, and A.-L. Barabási. The implications of human metabolic network topology for disease comorbidity. Proceedings of the National Academy of Sciences, 105(29):9880–9885, 2008. ISSN 0027-8424. doi: 10.1073/pnas.0802208105.

3. Hongzhong Lu, Feiran Li, Benjamín J. Sánchez, Zhengming Zhu, Gang Li, Iván Domenzain, Simonas Marcišauskas, Petre Mihail Anton, Dimitra Lappa, Christian Lieven, Moritz Emanuel Beber, Nikolaus Sonnenschein, Eduard J. Kerkhoven, and Jens Nielsen. A consensus S. cerevisiae metabolic model Yeast8 and its ecosystem for comprehensively probing cellular metabolism. Nature Communications, 10(1):3586, 2019. doi: 10.1038/s41467-019-11581-3.

4. Akhil Kumar, Lin Wang, Chiam Yu Ng, and Costas D. Maranas. Pathway design using de novo steps through uncharted biochemical spaces. Nature Communications, 9(1):184, 2018. doi: 10.1038/s41467-017-02362-x.

5. Jeanine A. Harrigan, Xavier Jacq, Niall M. Martin, and Stephen P. Jackson. Deubiquitylating enzymes and drug discovery: emerging opportunities. Nature Reviews Drug Discovery, 17(1):57–78, 2018. ISSN 1474-1776. doi: 10.1038/nrd.2017.152.

6. Sayada Reemsha Kazmi, Ren Jun, Myeong-Sang Yu, Chanjin Jung, and Dokyun Na. In silico approaches and tools for the prediction of drug metabolism and fate: A review. Computers in Biology and Medicine, 106:54–64, 2019. ISSN 0010-4825. doi: 10.1016/j.compbiomed.2019.01.008.

7. Sjoerd Slagman and Wolf-Dieter Fessner. Biocatalytic routes to anti-viral agents and their synthetic intermediates. Chemical Society Reviews, 50(3):1968–2009, 2020. ISSN 0306-0012. doi: 10.1039/d0cs00763c.

8. Roger A. Sheldon and John M. Woodley. Role of Biocatalysis in Sustainable Chemistry. Chemical Reviews, 118(2):801–838, 2018. ISSN 0009-2665. doi: 10.1021/acs.chemrev.7b00203.

9. Shuke Wu, Radka Snajdrova, Jeffrey C. Moore, Kai Baldenius, and Uwe T. Bornscheuer. Biocatalysis: Enzymatic Synthesis for Industrial Applications. Angewandte Chemie International Edition, 60 (1):88–119, 2021. ISSN 1433-7851. doi: 10.1002/anie.202006648.

10. Baudoin Delépine, Thomas Duigou, Pablo Carbonell, and Jean-Loup Faulon. RetroPath2.0: A retrosynthesis workflow for metabolic engineers. Metabolic Engineering, 45:158–170, 2018. ISSN 1096-7176. doi: 10.1016/j.ymben.2017.12.002.

11. Homa Mohammadi Peyhani, Jasmin Hafner, Anastasia Sveshnikova, Victor Viterbo, and Vassily Hatzimanikatis. Expanding biochemical knowledge and illuminating metabolic dark matter with ATLASx. Nature Communications, 13(1):1560, 2022. doi: 10.1038/s41467-022-29238-z.

12. Daniel Probst, Matteo Manica, Yves Gaetan Nana Teukam, Alessandro Castrogiovanni, Federico Paratore, and Teodoro Laino. Biocatalysed synthesis planning using data-driven learning. Nature Communications, 13(1):964, 2022. doi: 10.1038/s41467-022-28536-w.

13. David Kreutter, Philippe Schwaller, and Jean-Louis Reymond. Predicting enzymatic reactions with a molecular transformer. Chemical Science, 12(25):8648–8659, 2021. ISSN 2041-6520. doi: 10.1039/d1sc02362d.

14. Peter D. Karp, Daniel Weaver, and Mario Latendresse. How accurate is automated gap filling of metabolic models? BMC Systems Biology, 12(1):73, 2018. doi: 10.1186/s12918-018-0593-7.

15. Daniel Lowe. Chemical reactions from us patents (1976-sep2016), 2017.

16. Alex Bateman, Maria-Jesus Martin, Sandra Orchard, Michele Magrane, Emanuele Alpi, Benoit Bely, Mark Bingley, Ramona Britto, Borisas Bursteinas, Gianluca Busiello, Hema Bye-A-Jee, Alan Da Silva, Maurizio De Giorgi, Tunca Dogan, Leyla Garcia Castro, Penelope Garmiri, George Georghiou, Daniel Gonzales, Leonardo Gonzales, Emma Hatton-Ellis, Alexandr Ignatchenko, Rizwan Ishtiaq, Petteri Jokinen, Vishal Joshi, Dushyanth Jyothi, Rodrigo Lopez, Jie Luo, Yvonne Lussi, Alistair MacDougall, Fabio Madeira, Mahdi Mahmoudy, Manuela Menchi, Andrew Nightingale, Joseph Onwubiko, Barbara Palka, Klemens Pichler, Sangya Pundir, Guoying Qi, Shriya Raj, Alexandre Renaux, Milagros Rodriguez Lopez, Rabie Saidi, Tony Sawford, Aleksandra Shypitsyna, Elena Speretta, Edward Turner, Nidhi Tyagi, Preethi Vasudev, Vladimir Volynkin, Tony Wardell, Kate Warner, Xavier Watkins, Rossana Zaru, Hermann Zellner, Alan Bridge, Ioannis Xenarios, Sylvain Poux, Nicole Redaschi, Lucila Aimo, Ghislaine Argoud-Puy, Andrea Auchincloss, Kristian Axelsen, Parit Bansal, Delphine Baratin, Marie-Claude Blatter, Jerven Bolleman, Emmanuel Boutet, Lionel Breuza, Cristina Casals-Casas, Edouard de Castro, Elisabeth Coudert, Beatrice Cuche, Mikael Doche, Dolnide Dornevil, Anne Estreicher, Livia Famiglietti, Marc Feuermann, Elisabeth Gasteiger, Sebastien Gehant, Vivienne Gerritsen, Arnaud Gos, Nadine Gruaz, Ursula Hinz, Chantal Hulo, Nevila Hyka-Nouspikel, Florence Jungo, Guillaume Keller, Arnaud Kerhornou, Vicente Lara, Philippe Lemercier, Damien Lieberherr, Thierry Lombardot, Xavier Martin, Patrick Masson, Anne Morgat, Teresa Batista Neto, Salvo Paesano, Ivo Pedruzzi, Sandrine Pilbout, Monica Pozzato, Manuela Pruess, Catherine Rivoire, Christian Sigrist, Karin Sonesson, Andre Stutz, Shyamala Sundaram, Michael Tognolli, Laure Verbregue, Cathy H Wu, Cecilia N Arighi, Leslie Arminski, Chuming Chen, Yongxing Chen, Julie Cowart, John S Garavelli, Hongzhan Huang, Kati Laiho, Peter McGarvey, Darren A Natale, Karen Ross, C R Vinayaka, Qinghua Wang, Yuqi Wang, Lai-Su Yeh, and Jian Zhang. UniProt: a worldwide hub of protein knowledge. Nucleic Acids Research, 47(Database issue):gky1049.#x2013;, 2018. ISSN 1362-4962. doi: 10.1093/nar/gky1049.

17. Parit Bansal, Anne Morgat, Kristian B Axelsen, Venkatesh Muthukrishnan, Elisabeth Coudert, Lucila Aimo, Nevila Hyka-Nouspikel, Elisabeth Gasteiger, Arnaud Kerhornou, Teresa Batista Neto, Monica Pozzato, Marie-Claude Blatter, Alex Ignatchenko, Nicole Redaschi, and Alan Bridge. Rhea, the reaction knowledgebase in 2022. Nucleic acids research, 50(D1):D693–D700, 2021. ISSN 0305-1048. doi: 10.1093/nar/gkab1016.

18. Andrew G. McDonald, Sinéad Boyce, and Keith F. Tipton. ExplorEnz: the primary source of the IUBMB enzyme list. Nucleic Acids Research, 37(Suppl_1):D593–D597, 2009. ISSN 0305-1048. doi: 10.1093/nar/gkn582.

19. Amos Bairoch. The ENZYME database in 2000. Nucleic Acids Research, 28(1):304–305, 2000. ISSN 0305-1048. doi: 10.1093/nar/28.1.304.

20. Markus Meuwly. Machine Learning for Chemical Reactions. Chemical Reviews, 121(16):10218– 10239, 2021. ISSN 0009-2665. doi: 10.1021/acs.chemrev.1c00033.

21. Philippe Schwaller, Alain C. Vaucher, Ruben Laplaza, Charlotte Bunne, Andreas Krause, Clemence Corminboeuf, and Teodoro Laino. Machine intelligence for chemical reaction space. Wiley Interdisciplinary Reviews: Computational Molecular Science, 12(5), 2022. ISSN 1759-0876. doi: 10.1002/wcms.1604.

22. Zhenzhen Zou, Shuye Tian, Xin Gao, and Yu Li. mlDEEPre: Multi-Functional Enzyme Function Prediction With Hierarchical Multi-Label Deep Learning. Frontiers in Genetics, 9:714, 2019. ISSN 1664-8021. doi: 10.3389/fgene.2018.00714.

23. Vladimir Gligorijević, P. Douglas Renfrew, Tomasz Kosciolek, Julia Koehler Leman, Daniel Berenberg, Tommi Vatanen, Chris Chandler, Bryn C. Taylor, Ian M. Fisk, Hera Vlamakis, Ramnik J. Xavier, Rob Knight, Kyunghyun Cho, and Richard Bonneau. Structure-based protein function prediction using graph convolutional networks. Nature Communications, 12(1):3168, 2021. doi: 10.1038/s41467-021-23303-9.

24. Alperen Dalkiran, Ahmet Sureyya Rifaioglu, Maria Jesus Martin, Rengul Cetin-Atalay, Volkan Atalay, and Tunca Doğan. ECPred: a tool for the prediction of the enzymatic functions of protein sequences based on the EC nomenclature. BMC Bioinformatics, 19(1):334, 2018. doi: 10.1186/s12859-018-2368-y.

25. Jae Yong Ryu, Hyun Uk Kim, and Sang Yup Lee. Deep learning enables high-quality and highthroughput prediction of enzyme commission numbers. Proceedings of the National Academy of Sciences, 116(28):13996–14001, 2019. ISSN 0027-8424. doi: 10.1073/pnas.1821905116.

26. Masaaki Kotera, Yasushi Okuno, Masahiro Hattori, Susumu Goto, and Minoru Kanehisa. Computational Assignment of the EC Numbers for Genomic-Scale Analysis of Enzymatic Reactions. Journal of the American Chemical Society, 126(50):16487–16498, 2004. ISSN 0002-7863. doi: 10.1021/ja0466457.

27. Yoshihiro Yamanishi, Masahiro Hattori, Masaaki Kotera, Susumu Goto, and Minoru Kanehisa. Ezyme: predicting potential EC numbers from the chemical transformation pattern of substrate-product pairs. Bioinformatics, 25(12):i179–i186, 2009. ISSN 1367-4803. doi: 10.1093/bioinformatics/btp223.

28. Syed Asad Rahman, Sergio Martinez Cuesta, Nicholas Furnham, Gemma L. Holliday, and Janet M. Thornton. EC-BLAST: A Tool to Automatically Search and Compare Enzyme Reactions. Nature methods, 11(2):171–174, 2014. ISSN 1548-7091. doi: 10.1038/nmeth.2803.

29. Diogo A. R. S. Latino and João Aires-de Sousa. Assignment of EC Numbers to Enzymatic Reactions with MOLMAP Reaction Descriptors and Random Forests. Journal of Chemical Information and Modeling, 49(7):1839–1846, 2009. ISSN 1549-9596. doi: 10.1021/ci900104b.

30. Volker Egelhofer, Ida Schomburg, and Dietmar Schomburg. Automatic Assignment of EC Numbers. PLoS Computational Biology, 6(1):e1000661, 2010. ISSN 1553-734X. doi: 10.1371/journal.pcbi.1000661.

31. Qian-Nan Hu, Hui Zhu, Xiaobing Li, Manman Zhang, Zhe Deng, Xiaoyan Yang, and Zixin Deng. Assignment of EC Numbers to Enzymatic Reactions with Reaction Difference Fingerprints. PLoS ONE, 7(12):e52901, 2012. doi: 10.1371/journal.pone.0052901.

32. Pablo Carbonell, Jerry Wong, Neil Swainston, Eriko Takano, Nicholas J Turner, Nigel S Scrutton, Douglas B Kell, Rainer Breitling, and Jean-Loup Faulon. Selenzyme: enzyme selection tool for pathway design. Bioinformatics, 34(12):2153–2154, 2018. ISSN 1367-4803. doi: 10.1093/bioinformatics/bty065.

33. Yoshihiko Matsuta, Masahiro Ito, and Yukako Tohsato. ECOH: An Enzyme Commission number predictor using mutual information and a support vector machine. Bioinformatics, 29(3):365–372, 2013. ISSN 1367-4803. doi: 10.1093/bioinformatics/bts700.

34. Noushin Hadadi, Homa MohammadiPeyhani, Ljubisa Miskovic, Marianne Seijo, and Vassily Hatzimanikatis. Enzyme annotation for orphan and novel reactions using knowledge of substrate reactive sites. Proceedings of the National Academy of Sciences, 116(15):7298–7307, 2019. ISSN 0027-8424. doi: 10.1073/pnas.1818877116.

35. Joaquín Borrego-Díaz and Juan Galán-Páez. Explainable Artificial Intelligence in Data Science. Minds and Machines, 32(3):485–531, 2022. ISSN 0924-6495. doi: 10.1007/s11023-022-09603-z.

36. Tim Miller. Explanation in artificial intelligence: Insights from the social sciences. Artificial Intelligence, 267:1–38, 2019. ISSN 0004-3702. doi: 10.1016/j.artint.2018.07.007.

37. Gherman Novakovsky, Nick Dexter, Maxwell W. Libbrecht, Wyeth W. Wasserman, and Sara Mostafavi. Obtaining genetics insights from deep learning via explainable artificial intelligence. Nature Reviews Genetics, pages 1–13, 2022. ISSN 1471-0056. doi: 10.1038/s41576-022-00532-2.

38. Hui Wen Loh, Chui Ping Ooi, Silvia Seoni, Prabal Datta Barua, Filippo Molinari, and U Rajendra Acharya. Application of explainable artificial intelligence for healthcare: A systematic review of the last decade (2011–2022). Computer Methods and Programs in Biomedicine, 226:107161, 2022. ISSN 0169-2607. doi: 10.1016/j.cmpb.2022.107161.

39. Hassan Khosravi, Simon Buckingham Shum, Guanliang Chen, Cristina Conati, Yi-Shan Tsai, Judy Kay, Simon Knight, Roberto Martinez-Maldonado, Shazia Sadiq, and Dragan Gašević. Explainable Artificial Intelligence in education. Computers and Education: Artificial Intelligence, 3:100074, 2022. ISSN 2666-920X. doi: 10.1016/j.caeai.2022.100074.

40. Andrea Mastropietro, Giuseppe Pasculli, Christian Feldmann, Raquel Rodríguez-Pérez, and Jürgen Bajorath. EdgeSHAPer: Bond-centric Shapley value-based explanation method for graph neural networks. iScience, 25(10):105043, 2022. ISSN 2589-0042. doi: 10.1016/j.isci.2022.105043.

41. Henry Heberle, Linlin Zhao, Sebastian Schmidt, Thomas Wolf, and Julian Heinrich. XSMILES: interactive visualization for molecules, SMILES and XAI attribution scores. Journal of Cheminformatics, 15(1):2, 2023. ISSN 1758-2946. doi: 10.1186/s13321-022-00673-w.

42. Geemi P. Wellawatte, Aditi Seshadri, and Andrew D. White. Model agnostic generation of counterfactual explanations for molecules. Chemical Science, 13(13):3697–3705, 2022. ISSN 2041-6520. doi: 10.1039/d1sc05259d.

43. David Weininger. SMILES, a chemical language and information system. 1. Introduction to methodology and encoding rules. Journal of Chemical Information and Modeling, 28(1):31–36, 2 1988. ISSN 1549-9596. doi: 10.1021/ci00057a005.

44. Daniel Probst, Philippe Schwaller, and Jean-Louis Reymond. Reaction classification and yield prediction using the differential reaction fingerprint DRFP. Digital Discovery, 1(2):91–97, 2022. doi: 10.1039/d1dd00006c.

45. Antje Chang, Lisa Jeske, Sandra Ulbrich, Julia Hofmann, Julia Koblitz, Ida Schomburg, Meina Neumann-Schaal, Dieter Jahn, and Dietmar Schomburg. BRENDA, the ELIXIR core data resource in 2021: new developments and updates. Nucleic Acids Research, 49(D1):D498–D508, 2020. ISSN 0305-1048. doi: 10.1093/nar/gkaa1025.

46. David S Wishart, Carin Li, Ana Marcu, Hasan Badran, Allison Pon, Zachary Budinski, Jonas Patron, Debra Lipton, Xuan Cao, Eponine Oler, Krissa Li, Maïlys Paccoud, Chelsea Hong, An C Guo, Christopher Chan, William Wei, and Miguel Ramirez-Gaona. PathBank: a comprehensive pathway database for model organisms. Nucleic Acids Research, 48(D1):D470–D478, 2019. ISSN 0305-1048. doi: 10.1093/nar/gkz861.

47. Sébastien Moretti, Van Du T Tran, Florence Mehl, Mark Ibberson, and Marco Pagni. MetaNetX/MNXref: unified namespace for metabolites and biochemical reactions in the context of metabolic models. Nucleic Acids Research, 49(D1):gkaa992.#x2013;, 2020. ISSN 0305-1048. doi: 10.1093/nar/gkaa992.

48. Avanti Shrikumar, Peyton Greenside, and Anshul Kundaje. Learning Important Features Through Propagating Activation Differences. arXiv, 2017. doi: 10.48550/arxiv.1704.02685.

49. Scott Lundberg and Su-In Lee. A Unified Approach to Interpreting Model Predictions. arXiv, 2017. doi: 10.48550/arxiv.1705.07874.

50. Daniel Probst and Jean-Louis Reymond. A probabilistic molecular fingerprint for big data settings. Journal of Cheminformatics, 10(1):66, 2018. ISSN 1758-2946. doi: 10.1186/s13321-018-0321-8.

51. Daniel Probst and Jean-Louis Reymond. SmilesDrawer: Parsing and Drawing SMILES-Encoded Molecular Structures Using Client-Side JavaScript. Journal of Chemical Information and Modeling, 58(1):1–7, 2018. ISSN 1549-9596. doi: 10.1021/acs.jcim.7b00425.

52. Erik Bernhardsson. Annoy: approximate nearest neighbors in c++/python optimized for memory usage and loading/saving to disk. GitHub https://github.com/spotify/annoy, 2017.

